# Integrated cytometry with machine learning applied to high-content imaging of human kidney tissue for *in-situ* cell classification and neighborhood analysis

**DOI:** 10.1101/2021.12.27.474025

**Authors:** Seth Winfree, Andrew T. McNutt, Suraj Khochare, Tyler J. Borgard, Daria Barwinska, Angela R. Sabo, Michael J. Ferkowicz, James C. Williams, James E. Lingeman, Connor J Gulbronson, Katherine J. Kelly, Timothy A. Sutton, Pierre C. Dagher, Michael T. Eadon, Kenneth W. Dunn, Tarek M. El-Achkar

## Abstract

The human kidney is a complex organ with various cell types that are intricately organized to perform key physiological functions and maintain homeostasis. New imaging modalities such as mesoscale and highly multiplexed fluorescence microscopy are increasingly applied to human kidney tissue to create single cell resolution datasets that are both spatially large and multi-dimensional. These single cell resolution high-content imaging datasets have a great potential to uncover the complex spatial organization and cellular make-up of the human kidney. Tissue cytometry is a novel approach used for quantitative analysis of imaging data, but the scale and complexity of such datasets pose unique challenges for processing and analysis. We have developed the Volumetric Tissue Exploration and Analysis (VTEA) software, a unique tool that integrates image processing, segmentation and interactive cytometry analysis into a single framework on desktop computers. Supported by an extensible and open-source framework, VTEA’s integrated pipeline now includes enhanced analytical tools, such as machine learning, data visualization, and neighborhood analyses for hyperdimensional large-scale imaging datasets. These novel capabilities enable the analysis of mesoscale two and three-dimensional multiplexed human kidney imaging datasets (such as CODEX and 3D confocal multiplexed fluorescence imaging). We demonstrate the utility of this approach in identifying cell subtypes in the kidney based on labels, spatial association and their microenvironment or neighborhood membership. VTEA provides integrated and intuitive approach to decipher the cellular and spatial complexity of the human kidney and complement other transcriptomics and epigenetic efforts to define the landscape of kidney cell types.

## Introduction

Renal researchers increasingly appreciate the importance of characterizing the cellular niches of the kidney (cell types and subtypes, physiological state, neighborhood interactions) and how they are altered in kidney disease^1–4^. Imaging of kidney tissue specimens with single-cell resolution at a large scale is an attractive approach for uncovering cellular niches in their tissue context^5–7^. Such an approach has become more feasible because of the increased ability to collect mesoscale imaging datasets on accessible confocal microscopes, whole slide imagers and light-sheet microscopes. Furthermore, combining mesoscale imaging with highly multiplexed staining or labeling approaches allows for the capture, *in situ*, of hundreds-of-thousands cells that may be classified by many markers. The scale and depth of these data create a mounting challenge for timely quantitative and interpretable analysis. This is particularly important for kidney research, where biopsy-scale multiplexed imaging datasets of the human kidney are being collected and publicly released by large collaborative consortia such as the Kidney Precision Medicine Project (KPMP) and the Human BioMolecular Atlas Program (HuBMAP)^8,9^.

Tissue cytometry (TC) is a powerful approach for analyzing mesoscale fluorescence images with single-cell resolution^5,10–13^. Depending upon the imaging platform, datasets may be either 2D or 3D^10,14^. An important first step in TC is to survey all the cells by segmentation. This is often accomplished by using nuclei as fiduciaries for cells. Segmentation entails identifying regions of images as nuclei based on contrast provided by stains, registering each individual nucleus as an object, and identifying an associated cytoplasm spatially or with a specific marker. Features to describe these cells can be calculated on this segmentation. These features could be related to fluorescence intensities of markers within or around the nucleus (i.e. the cytoplasm) or based on the spatial coordinates or proximity relationships to these nuclei^10,13,14^. Multiplexing several markers in the same experiment enhances the richness of the imaging data, by providing specificity of cell types based on unique markers (or unique combination thereof) and generating spatial information based on the distribution of stains within the tissue^5,13–15^.

In a standard cytometry analysis, after cell segmentation, cells can be classified and quantitated by supervised approach using gating based on marker intensity, in which cells are defined according to threshold levels of marker fluorescence (gating). However, as the scale and complexity of multiplex mesoscale image volumes increase, such a manual approach becomes intractable and increasingly unlikely to be successful at uncovering the complex spatial organization and cellular niches of the kidney. Multiplexed mesoscale tissue cytometry thus requires tools supporting unsupervised analysis, ideally with machine learning, to characterize the cellular makeup of a tissue accurately and completely, to identify cellular niches and to map their neighborhoods and microenvironments^11,14,16–18^. Given the complexity of the interacting processes of segmentation, classification, quantification and neighborhood analysis, the ideal system should incorporate all these processes into a single, integrated analysis and visualization software package.

Since the initial description of <u>V</u>olumetric <u>T</u>issue <u>E</u>xploration and <u>A</u>nalysis (VTEA) as an open-source project, several excellent tools for tissue cytometry have been developed (**Table 1**). In this time VTEA (v0.5.2-v0.7) has been used in several projects involving single imaging fields to large mesoscale multi-fluorescence kidney image volumes to perform supervised cytometry analysis with gating^4,5,19–24^. However, machine learning, data visualization and analysis tools that are useful for multidimensional, big-data scale imaging data were not previously implemented in VTEA’s uniquely integrated pipeline. Here we describe VTEA 1.0 which specifically adds, 1) machine learning for clustering and dimensionality reduction to aid in automated classification of cell types, 2) neighborhood analysis to uncover cellular niches and 3) new data visualization tools to support discovery. To facilitate this growth of VTEA’s integrated approach, a SciJava framework was implemented^25,26^. VTEA now supports extensible image processing, segmentation, classification, visualization, and neighborhood analysis for processing on hundreds-of-thousands of cells and multi-gigabyte datasets with a fully integrated workflow. We demonstrate the utility of VTEA by identifying cell subtypes based on labels, spatial association and neighborhood membership using large scale 3D and 2D imaging data from kidney tissue.

**Table 1.**
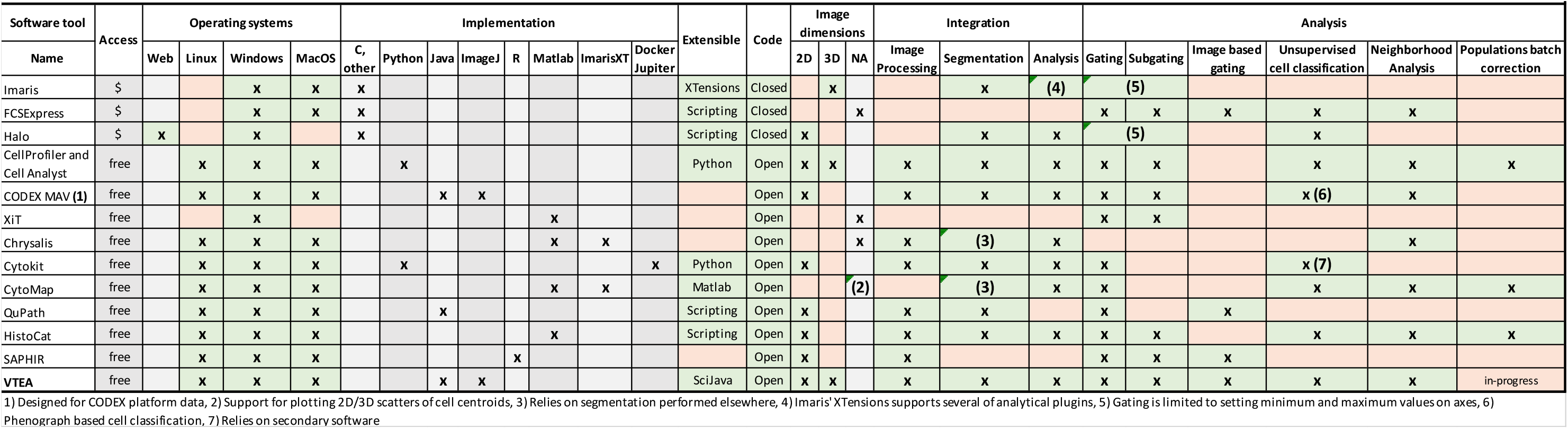
Tissue Cytometry Software.

## Methods

### Data acquisition and availability

Image data used in figures 3,4 and 5 or figure 1 and S2 was previously and separately analyzed in Woloshuk et al, 2021 or Winfree et al, 2017 ^10,23^. The analyses herein are wholly separate and independent of those analyses. The data present here is available upon request or, in-part, at https://vtea.wiki.

**Figure 1.**
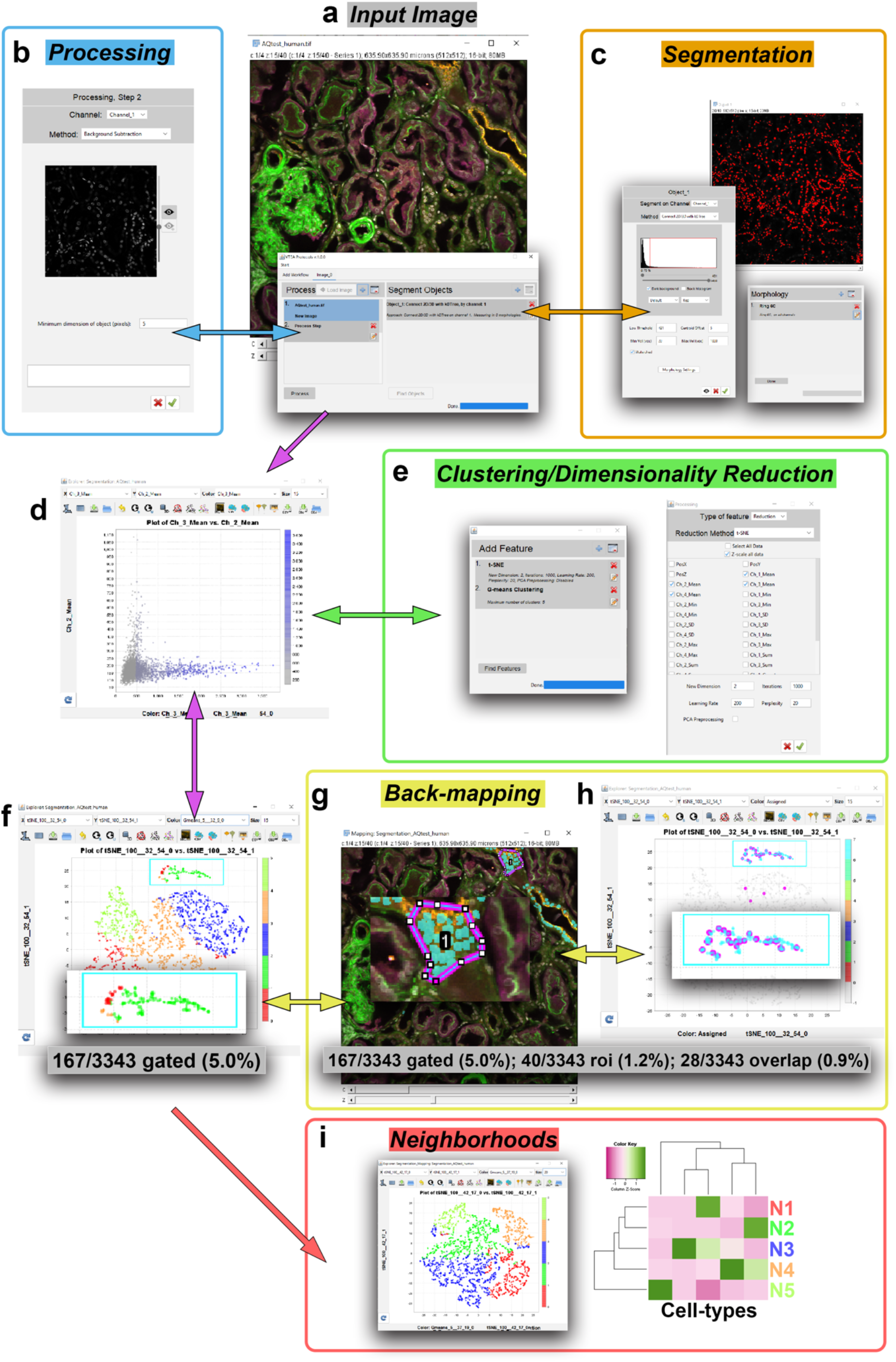
Integration of VTEA into a renal tissue cytometry workflow. VTEA provides an interactive and bidirectional workspace to integrate the image processing (Processing, blue, **b**), segmentation (Segmentation, orange, **c**), clustering and dimensionality reduction (green, **e**), mapping and back-mapping between the image and scatter plots (yellow, **g** and **h**) and generation of neighborhoods (Neighborhoods, red, **i**). Neighborhoods generated in VTEA can be clustered, back-mapped and additional levels of micro-environments (**e-i**).

### Tissue acquisition

Tissue was collected and processed under the Institutional Review Board at Indiana University approved protocols: 1906572234, for nephrectomy samples and 1010002261, for human biopsy samples.

### Tissue preparation

#### Multiplexed Immunofluorescence

Sectioning and staining for multiplexed staining were performed as described previously^23^. Briefly, 50 um sections of human nephrectomy tissue were cut from formaldehyde fixed (4.0% and stored in 0.25% in 1X PBS) with a vibratome (Leica Biosystems). Tissue sections were blocked in 10% normal donkey serum, 0.1% Triton X-100 in 1X phosphate buffered saline (PBS) and stained overnight with antibodies in block (**Table 3**)^7^. All secondaries were raised in donkey (ThermoFisher). In some cases, primary antibodies were directly conjugated including CD31, CD45, and Nestin conjugated with Dylight550, Alexa488 and Alexa647 (Thermofisher). After staining with DAPI for nuclei and in some cases with Alexa488-phalloidin for F-actin for 30 minutes, cells were washed in block, 1X PBS and mounted in Prolong Gold or Prolong Glass under #1.5 coverslips (ThermoFisher). Prolong Glass mounts were allowed to cure for 24-48 hours. As required coverslips were sealed with clear nail polish (Electron Microscopy Supply) and stored at 4°C before imaging.

#### Highly multiplexed immunofluorescence

Sectioning and staining for highly multiplexed immunofluorescence were performed as described previously^5,27^. Briefly, 50 um human nephrectomy tissue was cryosectioned and fixed in formaldehyde (4.0% and stored in 0.25% in 1X PBS). Blocking, staining was performed with six antibodies, secondaries as needed, DAPI and phalloidin as indicated (**Table 3**). Coverslips were mounted in Prolong Glass under #1.5 coverslips and cured for 24-48 hours. The eight markers were collected across 16 channels with four excitation wavelengths and spanning the visible spectrum and near infra-red.

### Confocal Microscopy

Multiplexed immunofluorescence specimens were imaged on either an FV1000 (Olympus) at 20x 0.75 NA oil immersion objective or a fully automated SP8 confocal with a 20x 0.75 NA Mimm objective as described previously^23^. Highly multiplexed fluorescence imaging was performed on the fully automated SP8 confocal and unmixed and stitched in either LASX(Leica) or FIJI^26^.

### CODEX

#### Antibody Conjugation and Validation

14 of the 23 antibodies used here were conjugated in-house using the protocol outlined by Akoya Biosciences (**Table 3**)^19,20^. To conjugate barcodes to antibodies, antibodies were reduced using a “Reduction Master Mix” (Akoya Biosciences) to which lyophilized barcodes resuspended in molecular biology grade water and “Conjugation Solution” (Akoya Biosciences) were added and incubated for 2 hours at room temperature. Labeled antibodies were purified from free-barcode in a 3-step wash and spin process and stored at 4°C. Successful conjugation was validated via SDS-PAGE gel electrophoresis as well as immunofluorescent staining of reference tissue followed by confocal microscopy.

#### Tissue Preparation

10-micron sections of human renal tissue embedded in OCT were cut onto poly-L-lysine coated coverslips. Sections were prepared as detailed by Akoya Biosciences and as described previously^7,14^. Tissue retrieval was conducted with a 3-step hydration process, followed by fixation with a PFA-containing solution. Following fixation, the coverslip mounted tissue was incubated overnight at 4°C with an antibody cocktail of 23 of the antibodies listed in **Table 3**. Tissues were washed, and post-fixed in 4% PFA for 15 minutes. **Imaging**. Antibodies were imaged cyclically using the CODEX system from Akoya Biosciences and a Keyence BZ-X810 slide scanning microscope fitted with a 20x air objective (0.75 NA). Images were processed using the CODEX Processor (Akoya Biosciences) and images exported for analysis with VTEA.

### Software design, development, and distribution

Volumetric Tissue Exploration and Analysis (v1.0) was developed in Java, with SQL, R-script and Python, using the integrated development environment Netbeans (Apache) using a maven build scheme. Major application program interfaces (APIs) used in VTEA include SciJava (v.30.0.0), ImageJ (v.1.53f), h2 (v. 1.4.198), SMILE (v.1.5.3), Renjin (v3.5-beta76) and JFreeChart (v. 1.5.0). The github release tag @cdfbd46 can be used to perform all the analyses presented here. VTEA v1.0, bleeding-edge and archival versions, can be downloaded and built from source-code using a maven build scheme, https://github.com/icbm-iupui/volumetric-tissue-exploration-analysis. Stable releases can be installed in FIJI by using the FIJI updater and selecting the “Volumetric Tissue Exploration and Analysis” update site. General description, analysis vignettes with demonstration data and development plans can be found at https://www.vtea.wiki and https://imagej.net/plugins/vtea.

### Computers used in analysis

Image data was analyzed VTEA on a Macbook laptop (mCorei5, 8 GB RAM, 2016), a Lenovo P51 (Xeon quad-core, 64 GB RAM) or an 8-core custom-built workstation (Xeon 8-core, 256 GB RAM).

### Figure preparation

All images were generated in ImageJ/FIJI and plots (scatter, violin, heatmaps), gated cell overlays and segmentation maps were created by VTEA. Photoshop (Adobe) was used for final annotation and assembly of panels. Scales were set by the microscopy platform and annotated in ImageJ/FIJI. All intensity changes are linear unless otherwise noted.

## Results

### VTEA, an integrated tool for image processing, segmentation, and interactive cytometry analysis

VTEA has a fully integrated analytical pipeline for image processing, segmentation, and cytometry in one software framework (**Figure 1**). Using the standard approach of VTEA, segmented cells can be analyzed in a 2D scatter plot, where gating can be applied based on marker intensity to identify and count specific cells of interest. Gated cells are visualized in the image volume, which allows for visual validation of the gating strategy and facilitate biological interpretation (**Figure 1f**). Furthermore, using ImageJ/FIJIs region-of-interest(roi) selection tool allows quantitation and visualization of select cells in the image within the scatter plot (**Figure 1g** and **h**). This back-and-forth interaction between the image and analytical space allows for fine-tuning of the analysis and interactive exploration. VTEA has also a unique bidirectional workflow, whereby a user can make upstream adjustments in image processing and segmentation and test the effect of these adjustments on the cytometry output (**Figure 1**, double headed arrows).

### Building an extensible framework for an integrated tissue cytometry (TC) analysis in VTEA

To facilitate on-going improvements to VTEA’s integrated pipeline, a SciJava based plugin infrastructure was implemented. The SciJava extensible framework uses interfaces implemented by super-classes that are extended by concrete implementations which include runtime annotations for indexing of available classes or plugins(**Table 2**). Using this extensible framework, new functionality was incorporated into VTEA including functionality from SMILE, Renjin, and ImageJ/FIJI(**Figure S1A**)^26,28,29^. To improve data handling the SQL based Java database h2 was also implemented (**Figure S1A**)^30^. Furthermore, additional functionality has been added to VTEA including but not limited to: 1) compatibility with other segmentation and analysis tools, 2) archiving and sharing data 3) generating multiclass training dataset for use in image classification (**Figure S1B**)^23^.

**Table 2.** SciJava Object Structures and Functionality.

| Functionality | Interface | Superclass | Comment |
| --- | --- | --- | --- |
| Factories, processors | Processor | AbstractProcessor.class | Defines a suite of factories for processing data and generating a result. |
| Image Processing | ImageProcessing | AbstractImageProcessing.class | Defines image processing steps to perform on imaging data and returns a processed copy. |
| Segmentation | Segmentation | AbstractSegmentation.class | Defines a segmentation approach and returns segmented objects. |
| Measured morphologies | Morphology | AbstractMorphology.class | Defines regions associated with segmented nuclei in which to measure intensity. |
| Measurements to make on segmented objects | Measurement | AbstractMeasurement.class | Defines the maths or statistics to perform on the pixel/voxel values associated with segmented objects. |
| Measurements to make on neighborhoods | NeighborhoodMeasurement | AbstractNeighborhoodMeasurement.class | Defines the measurements that are done on neighborhood class including summary statistics of constituents. |
| Features derived from unsupervised analysis | Feature | AbstractFeature.class | Defines an algorithm to perform unsupervised analysis on segmented cells based on measured features including AbstractMeasurements.class and AbstractNeighborhoodMeasurement.class. |
| Look up tables | LUT | AbstractLUT.class | Defines a LUT to be used in plot overlays and static plots generated by VTEA. |
| Static plots | Plot | AbstractPlot.class | Defines a plotting routine that returns an image file, e.g. .png. |

### VTEA with unsupervised machine learning improves the accuracy and resolution of cell classification in kidney image volumes

Manual gating has limitations in accuracy and resolution of cell classification and may underperform in the setting of large multi-dimensional datasets. The use of scatterplots, gates and sub-gating is a common strategy for analyzing cytometry data. Gating frequently requires the use of expert opinion and operator-dependent thresholding of cells that are positive or negative for a marker of interest (**Figure 2A**). For instance, in 3D confocal imaging data of human renal cortex, detecting aquaporin-2 (AQP2) positive cells is expected to identify the collecting duct (CD). However, in a standard VTEA analytical 2D scatter plot, the population of AQP2-positive cells may not be accurately nor rigorously identified by manual gating (**Figure2A**, lower right plot). Using one of VTEA’s newly implemented clustering algorithms, agglomerative hierarchical clustering with Ward linkage, using fluorescence intensities, the AQP2-positive population was readily clustered (**Figure 1e, Figure2B**, lower right). Importantly, this added functionality to VTEA also identified additional AQP2-positive cells not initially included in the expert drawn gate, and clearly a part of the CD epithelium when mapped back to the imaging volume improving the classification of putative AQP2-positive CD cells (**Figure 2C**).

**Figure 2.**
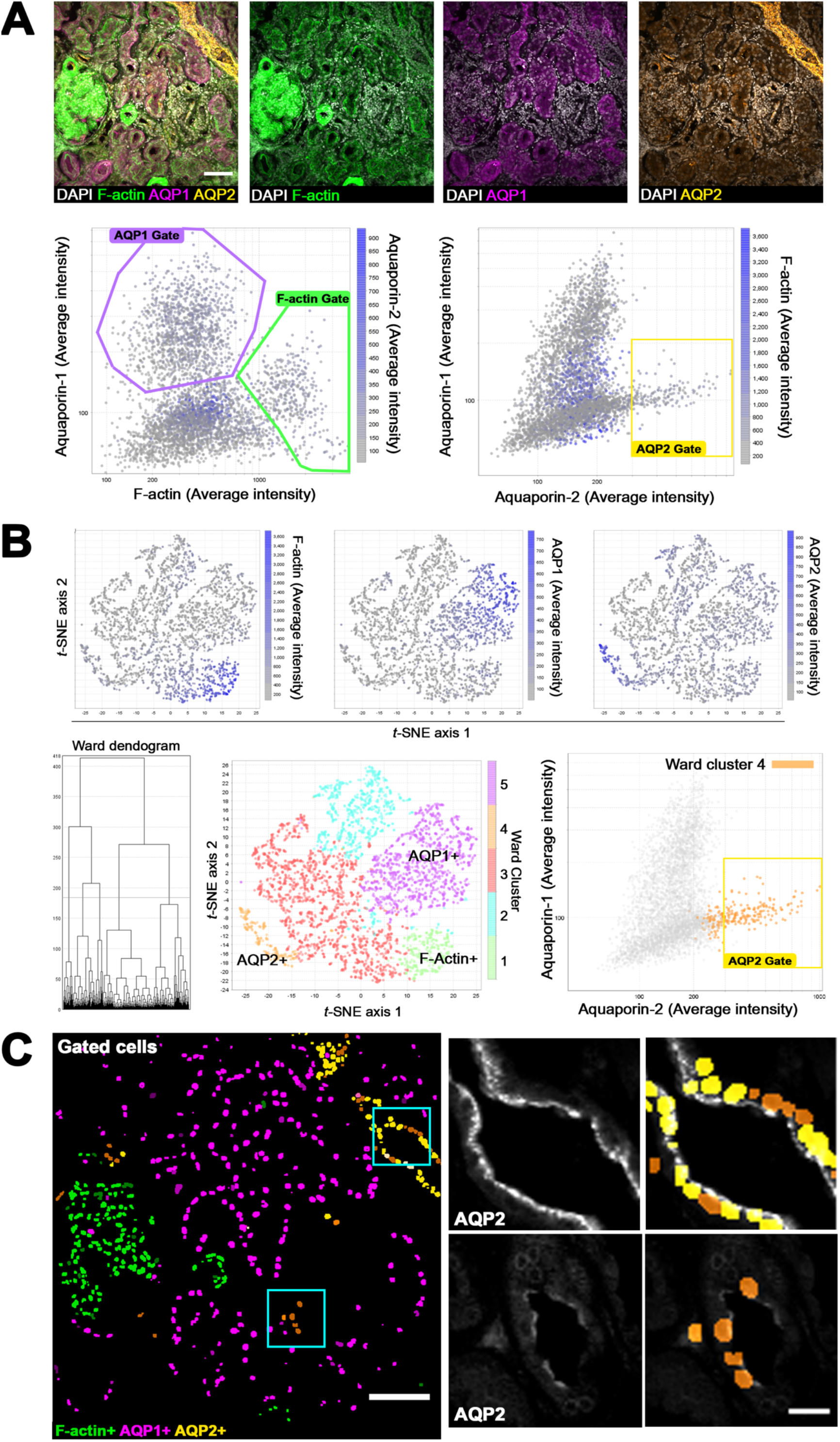
Improved classification of the kidney epithelium with semi-automated machine learning. 50-micron thick human kidney tissue was stained for proximal tubule and collecting duct markers aquaporin-1 (AQP1) and aquaporin-2(AQP2) and counterstained for nuclei (DAPI) and filamentous actin (F-actin, phalloidin) and imaged by confocal fluorescence microscopy. **A**. Supervised analysis of a volume of human kidney cortex in 3D (top panels) in VTEA with user defined gates based on the intensity of AQP1, AQP2, and F-actin intensities where the AQP2 gate was selected to avoid cells with high F-actin intensity per a lookup-table (LUT, lower panels). Scale bar = 100 um. **B**. Unsupervised dimensionality reduction and clustering in VTEA of average and standard deviation of cell associated intensities of DAPI, phalloidin, AQP1 and AQP2 partitions the cells into discrete groups. Intensity of F-actin, AQP1 and AQP2 is plotted as the LUT in *t*-SNE space (top panels) and Elbow Plot for estimation of k from selected feature space (k=5, vertical red line), Agglomerative hierarchical clustering with Ward-linkage (k=5, red line on dendogram) subdivides the cells into clusters that reflect the marker intensity (bottom panels). **C**. Changes in cell classification visualized in VTEA demonstrate an increase in the number of each group of cells with the unsupervised approach in (B). Cells classified in (A) are in yellow and cells in orange were added to the classification with Ward Hierarchical clustering (cluster 4, **B**). Scale bars = 100 and 20 um.

To improve the resolution of classified AQP2-positive populations, non-proximal epithelial cells (AQP1-, F-actin-negative cells) in the renal cortex (clusters 2,3 and 4, **Figure S2A and S2Ba**) were subgated in VTEA. Using fluorescence intensities, the segmented cells were re-clustered and mapped to *t-*SNE space with VTEA. Ward clustering uncovered three AQP2-positive clusters (**Figure S2Bb**) that mapped to distinct tubular epithelium. The two AQP2-positive clusters with highest intensity (clusters 1 and 2 **Figure S2Bc**) mapped predominantly to collecting duct cells and putative distal epithelial cells (verified by morphology), which could be consistent with APQ2-positive progenitor cells described previously^31^. The AQP2-positive with low intensity mapped to the distal tubular epithelium, suggesting a subpopulation of cells in the distal nephron with a low level of AQP2 (**Figure S2Ad** and **S2Bc**). The increased resolution and distribution of AQP2 expression was only apparent after using VTEA to 1) sub-gate and re-cluster putative APQ2-positive cells and 2) confirm these new populations with both mapping the gated cells to the image and selecting the tubules with an ROI in the image to map to the scatterplot (**Figure S1Ae** and **S1Bc**). Using VTEA’s unsupervised analysis, subgating and bi-directional mapping increased the resolution of cell classification which could not be performed using a traditional supervised approach (**Figure 1A**).

### Separation of spatially overlapping cell populations in 3D image volumes with unsupervised machine-learning and determination of glomerular census at large scale with VTEA

Using a manual gating strategy with standard 2D scatter plots may poorly discriminate spatially congested cells (for example within glomeruli), where staining from one cell can confound the classification of neighboring cells. As shown in **Figure 3A**, Gating Nestin-positive podocytes cells successfully in a standard scatter plot with VTEA is not feasible, because the analysis is confounded by the close association of Nestin-positive cells with CD31-positive glomerular endothelial cells and CD45-positive immune cells. Using dimensionality reduction with *t*-SNE, feature plots and unsupervised agglomerative hierarchical clustering with Ward-linkage as implemented in VTEA, Nestin-positive podocytes can be isolated from CD31-positive endothelial and CD45-positive immune cells (**Figure 3B**). Such an analytical approach can also be applied at a large scale, where unsupervised analysis can identify immune, endothelial and podocyte cells for an entire section of kidney tissue (**Figure 4A-B**). Glomerular cell census in such tissue can be readily acquired by applying spatial regions-of-interest on the image volume (**Figure 4D**). Cells in the glomeruli are counted and classified as podocytes, endothelial, immune or other (likely mesangial). Furthermore, the 3D volumes collected allow for volumetric cell densities to be calculated for glomeruli (**Figure 4D**).

**Figure 3.**
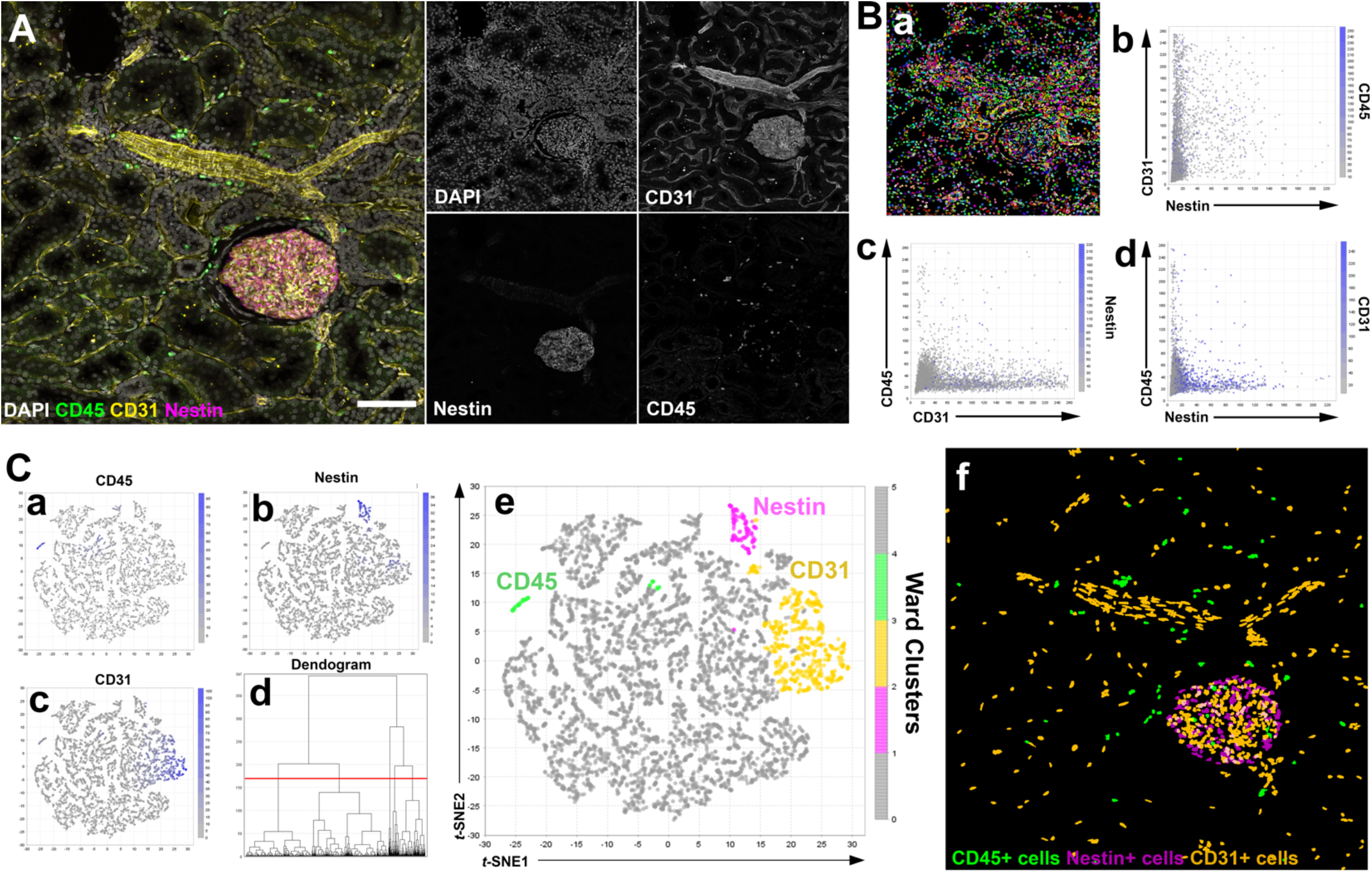
Separating masked populations of podocytes, leukocytes and endothelium by unsupervised analysis of cell image volumes in VTEA. **A**. Human nephrectomy tissue was fixed in and stained with DAPI and with antibodies against CD31 (yellow), CD45 (green) and Nestin (magenta) and imaged by confocal microscopy. Separated channels are given at right. All images are maximum intensity Z-projections, scale bar = 100 um. **B**. Nuclei segmented as fiduciaries of cell with connected components in 3D, ‘Connect 3D’, and associated intensities of CD31, CD45 and Nestin were measured. **Ba**. map of all segmented cells uniquely colorized. **Bb-d** Scatterplots of CD31, CD45 and Nestin demonstrate overlap in Nestin and CD31 signal in 2D plots. **C**. Unsupervised analysis of cell and marker intensity uncovers closely associated Nestin and CD31-positive cells. **Ca-Cc**. Mapping of cell associated intensity (every dot is a cell) to *t*-SNE projection calculated from the mean intensity of DAPI, CD31, CD45 and Nestin. **Cd**. Ward clustering dendogram using the same mean intensities with major bifurcation between labeled and unlabeled and at least 3 subclusters of labeled cells at k=5 (red line). **Ce**. Ward clusters mapped to *t-*SNE projection. Putative cell types are colored and labeled. **Cf**. Colorizing of segmented pixels for cells within CD45-positive, Nestin-positive and CD31-positive classes as given in **Ce**.

**Figure 4.**
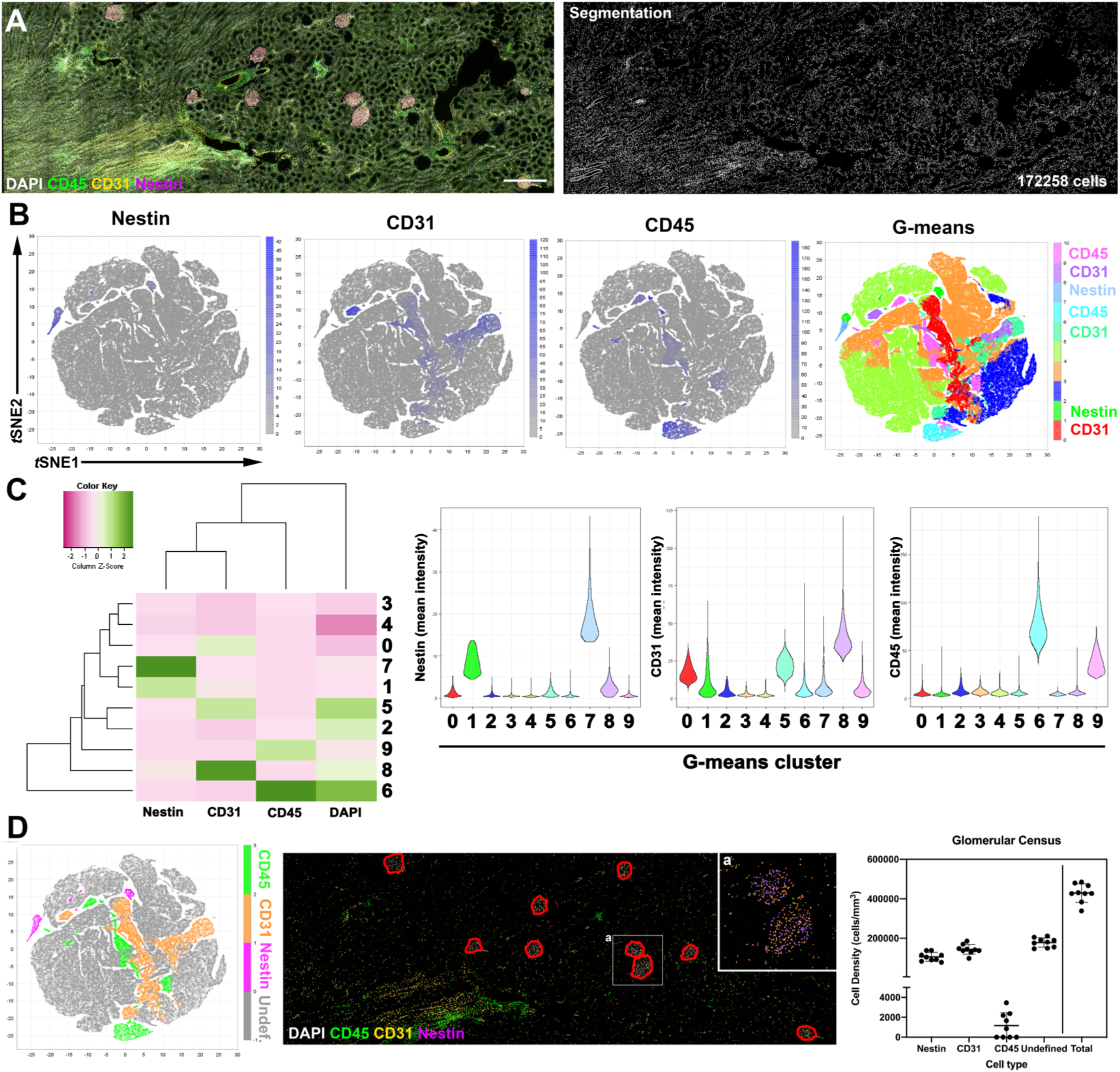
Automated glomerular census in mesoscale 3D tissue cytometry with machine learning classification and spatial analysis in VTEA. **A**. Human nephrectomy tissue was fixed in and stained with DAPI (gray) and with antibodies against CD31 (yellow), CD45 (green) and Nestin (magenta) and imaged by confocal microscopy. Scale bar = 500 um **B**. The whole 3D volume was analyzed, and 172,258 cells were segmented and processed by VTEA. Unsupervised analysis, G-means clustering and *t*-SNE mapping identified CD45-positive, CD31-positive, and Nestin-positive cells automatically. **C**. The cluster identities were confirmed by the plots generated by VTEA including average mean fluorescence intensity of each class, (heatmap, left panel) or by the distribution of mean fluorescence intensities for each cell (violin plots, right panels). **D**. Unsupervised classifications were used in combination (left panel) with regions-of-interest (ROIs) drawn with ImageJ during analysis to perform a glomerular census of the endothelium (CD31), leukocytes (CD45) and podocytes (Nestin). The census is presented as density of cells per mm^3^ (right panels).

### Assessing infiltrating immune cell subtypes in human kidney based on spatial neighborhoods

Focusing on the CD45-positive immune cells from Figure 3 and applying VTEA’s unsupervised analysis, we uncovered 2 populations of CD45-positive immune cells, which separated based on their nuclear staining and CD45 intensity (**Figure 5A and B**). We asked if these two subpopulations could distribute in unique spatial niches. Using the cell-centric neighborhood analysis function and data visualization features within VTEA, we defined the spatial neighborhoods of all labeled cells, including the two types of CD45-positive immune cells defined by CD45 and DAPI intensity where CD45-pop1 cells had higher intensities of CD45 and DAPI than CD45-pop2 (**Figure 5e**). Most neighborhoods with immune cells had a mixture of the two CD45-positive populations (**Figure 5Ca**,**5Cb** and **5Cg** neighborhoods 7, 8 and 9). However, a unique immune neighborhood enriched with endothelial cells (i.e., close to vessels) had a preponderance of CD45-pop1 (**Figure 5Ce** and **5Cg** Neighborhood 4). Another neighborhood with the lowest fraction of endothelial cells (i.e., distant from vessels) had a mostly of CD45-pop2 cells (**Figure 5Cf** and **5Cg** neighborhood 10). These findings demonstrate that unsupervised clustering with neighborhood analysis in VTEA can uncover a variety of cell niches based on spatial relationships that may have biological relevance.

**Figure 5.**
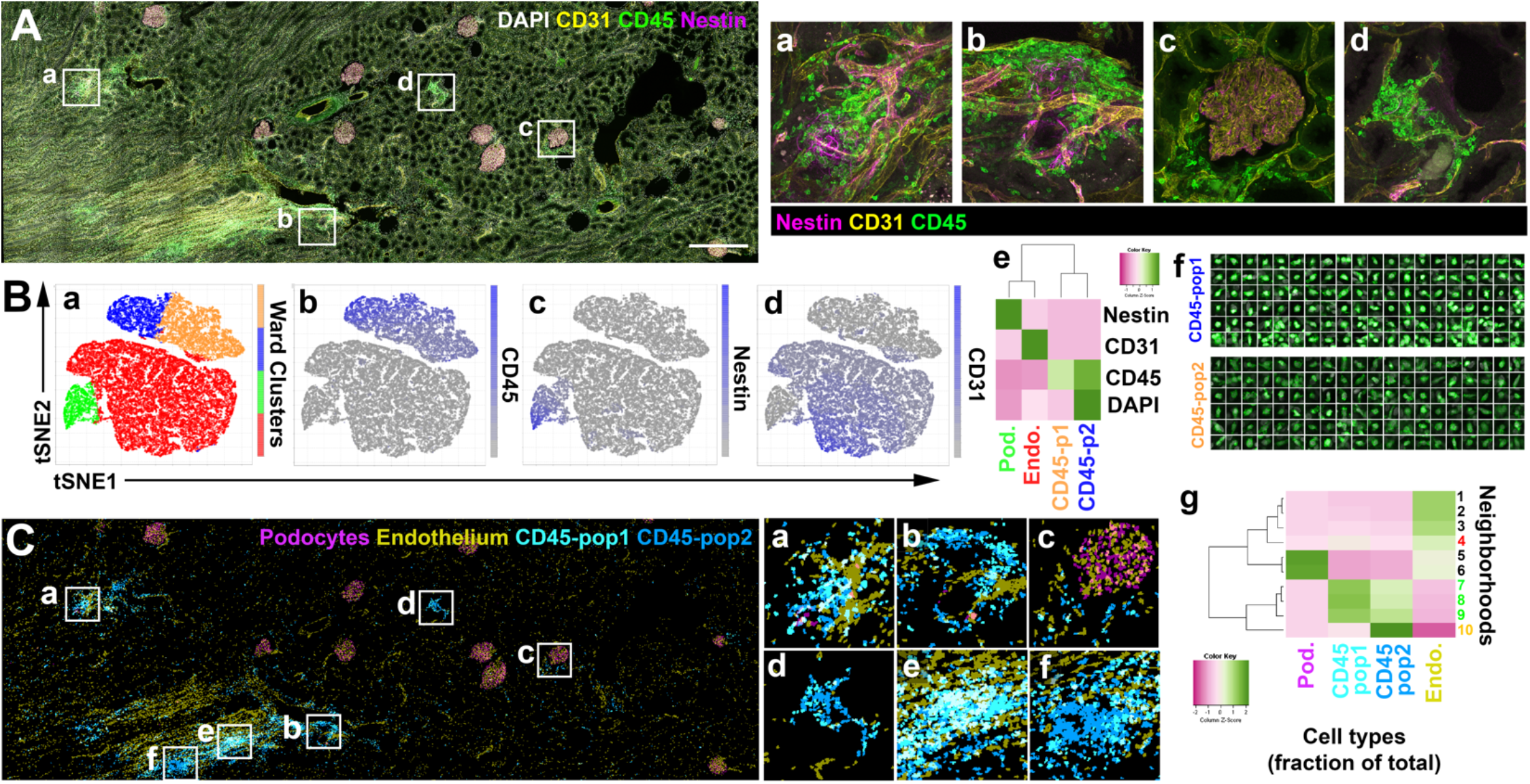
Using VTEA to classify leukocytes based on association with vascular endothelium. Using the image volume collected for Figure 4, spatially defined neighborhoods with CD45-positive cells were generated in VTEA. **A**. Mesoscale maximum z-projection of human nephrectomy tissue stained with DAPI (gray) and with antibodies against [CD31 (yellow), CD45 (green) and Nestin (magenta) and presented as separate channels in gray. CD45-positive leukocytes localize in clusters that associate with CD31-positive structures including glomeruli and blood vessels (**Aa-Ad**). Scale bar = 500 um **B**. CD45-positive leukocytes and CD31+ endothelium was subgated from all cells (Figure 4B) and reclustered for k=5. **Ba-Be**, feature plots and VTEA generated heatmaps of mean signal intensity for CD45, Nestin and CD31 uncovers podocytes (Pod., green), Endothelium (Endo., red) and two putative populations of CD45-positive cells based on DAPI and CD45 intensity (p1 or pop1 vs. p2 or pop2, orange and blue respectively). **Bf**. Using cell-wise export with VTEA ground truth generation routine, 100 cells sampled from the two CD45-positive populations. **Ca-Cb**. Maximum projections of mapped CD45-p1 and CD45-p2 in the image volume identifies tissue regions of mixed, and uniquely CD45-p2 cells. **Cg**, neighborhood analysis, for every classified cell within r = 25 um and unsupervised clustering of neighborhoods based on cellular census, demonstrates CD45-p1 are found in neighborhoods with (N4, red) and without endothelium (Endo, N7-9, green e.g., **Ca** and **Ce**) in neighborhoods with CD45-p2. CD45-p2 are also found alone without CD45-p1 nor endothelium (N10, orange, e.g., **Cd** and **Cf**).

### Using VTEA for classification of cell-types in 3D spectral confocal images of human kidney

The use of multispectral confocal imaging and linear unmixing of fluorophores enables large-scale 3D imaging with 8 multiplexed probes (**Figure 6A**). This approach allows the survey of tens-to hundreds-of-thousands of cells *in situ* within kidney tissue. Identifying major cells subtypes and the less abundant immune cells can be difficult using traditional gating, particularly when the emission spectra from fluorophores are overlapping. Using unsupervised machine learning, including clustering (**Figure 6Ba**-**6Bc**), and visualizing mean fluorescence intensity distributions of the cell subtypes in VTEA (**Figure 6Bd**), we can readily classify both epithelial and immune cells in an unbiased manner (**Figure 6B**).

**Figure 6.**
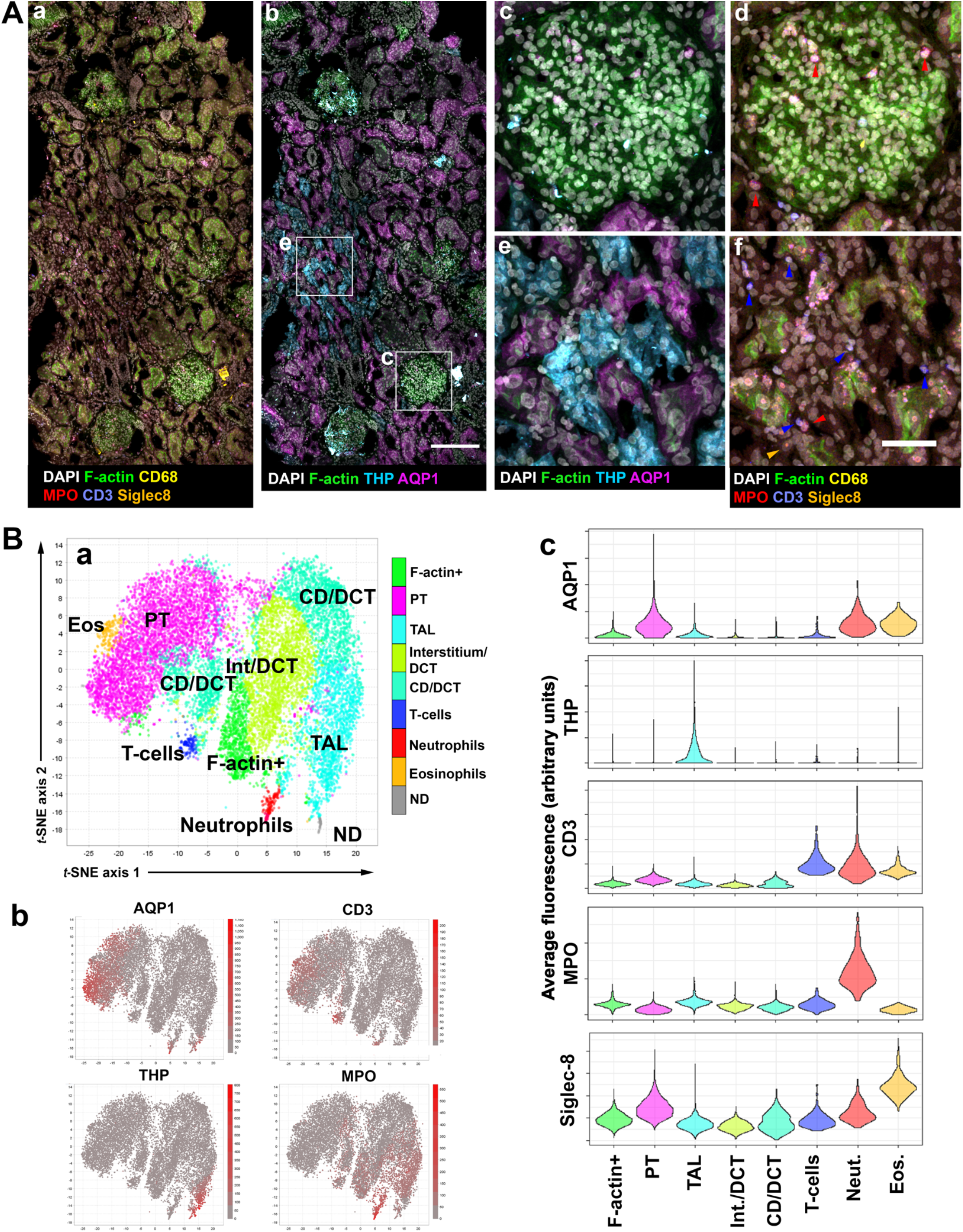
Uncovering cell types in 3D highly multiplexed data with VTEA. Human nephrectomy tissue was fixed and stained for 8 markers and imaged by 3D spectral confocal microscopy. Using reference spectra, 8 channels were unmixed and analyzed by VTEA. **A**. Mesoscale, maximum projections of subsets of channels highlighting immune or tubular epithelium (**Aa** and **Ab**), with insets as given in **Ab**. Arrowheads indicate neutrophils (red), T-cells (blue) and eosinophils (yellow). Scale bars are 200 um and 50 um. **B**. Unsupervised analysis of all cells with 8 marker channels separates the major cell types expected in the renal cortex. Mixed classes of Interstitium (Int) and distal nephron including distal convoluted tubule (DCT) or collecting ducts (CD) are low for all markers (**Bb** and **Bc**). A small group of THP high cells were annotated as not defined (ND, **Ba**). Granulocytes, eosinophils and neutrophils, are abbreviated as Neut and Eos in **Bc**.

### Applying VTEA’s pipeline on highly multiplexed CODEX data of the human kidney to uncover cell sub-types and biologically relevant cell neighborhoods

Human cortical biopsy underwent CODEX imaging with 23 markers (**Figure 7A, Table 3**) and analyzed with VTEA to perform cytometry, cell-classification, and neighborhood analysis (**Figure 7B-E**). Unsupervised hierarchical clustering of the 11,355 segmented cells identified the major cell types in the kidney (**Figure 7B**). Using subgating and subclustering, additional novel cell state phenotypes were identified including PROM1-positive (CD133) thick ascending limb and proximal tubule cells (**Figure 7B**, and **S3D**) and CD68-positive dendritic cells, CD68-positive putative macrophages, and CD68-positive epithelial cells (**Figure 7C** and **S3C-E**). To determine if specific cellular microenvironments were present in the tissue, VTEA’s neighborhood analysis function (**Figure 1i**) was used to tabulate neighborhoods and revealed, after clustering and dimensionality reduction, unique neighborhoods with associations between tubular epithelial cell subtypes and immune cells, such as the association of PROM1-positive TAL and PROM1-positive PT cells with low levels of CD68-positive macrophages (**Figure 7D** and **E**, N4 and N5 respectively) and an association of CD90-positive proximal tubules with DC and T-cells (**Figure 7E**, N6).

**Table 3.** Antibodies.

| Antibody | Clone | Supplier | Catalog Number | CODEX | Confocal |
| --- | --- | --- | --- | --- | --- |
| CD68 | PG-M1 | Agilent-Dako | M0876 |  | x |
| CD31 | JC/70A | NovusBio | NB600-562 |  | x |
| CD3 | UCHT1 | BD/Akoya | 557706/4350008 | x | x |
| Nestin | 10C2 | NovusBio | NB300-266 | x | x |
| Uromodulin | Polyclonal | R&D | AF5144 | x | x |
| AQP1 | 1/22 | Santa Cruz | sc-32737-X | x | x |
| MPO | Polyclonal | Abcam | ab9535 | x | x |
| CD45 | HI30 | Biolegend/Akoya | 304002/4150003 | x | x |
| LRP2 | Polyclonal | Abcam | ab76969 | x | x |
| Podocalyxin | Polyclonal | R&D | AF1658 | x |  |
| Calbindin | CB-955 | Abcam | ab82812 | x |  |
| Cytokeratin8 | TS1 | NovusBio | NBP2-34501-0.1mg | x |  |
| a-sma | 1A4 | Invitrogen | 14-9760-82 | x |  |
| PROM1 | AC133 | Miltenyi Biotec | 130-090-422 | x |  |
| CD68 | KP1 | ThermoFisher | 14-0688-82 | x |  |
| IGFBP7 | Polyclonal | Acris/Origene | AP01109PU-S | x |  |
| p- c-Jun | D47G9 | Cell Signalling | 3270BF | x |  |
| CD4 | SK3 | Akoya | 4350010 | x |  |
| CD11c | S-HCL-3 | Akoya | 4350012 | x |  |
| CD31 | WM59 | Akoya | 4250009 | x |  |
| CD45RO | UCHL1 | Akoya | 4250023 | x |  |
| HLA-DR | L243 | Akoya | 4250006 | x |  |
| CD90 | SE10 | Akoya | 4150021 | x |  |
| E-cadherin | 4A2C7 | Akoya | 4250021 | x |  |
| b-catenin | 12F7 | Akoya | 4450036 | x |  |

**Figure 7.**
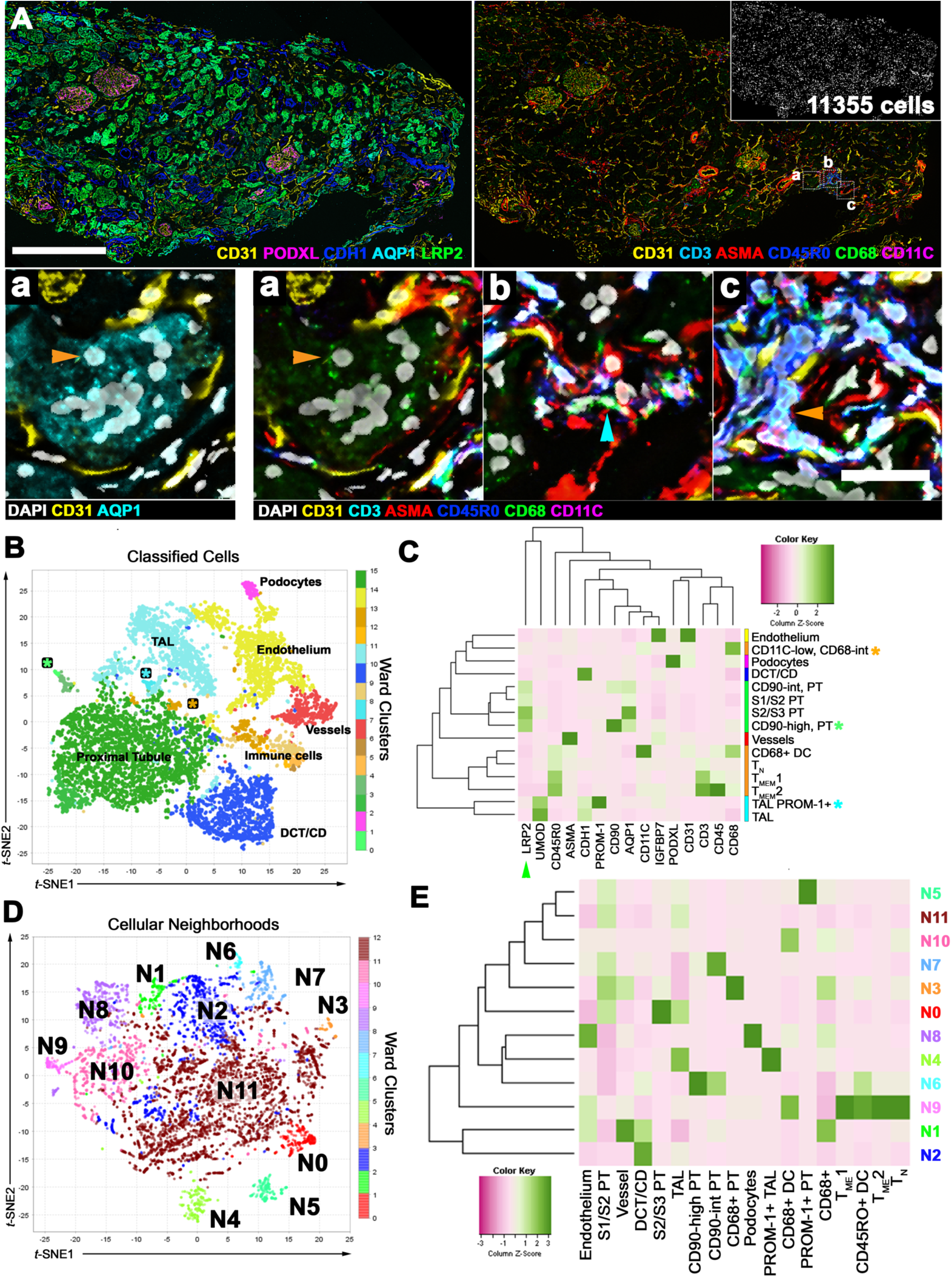
Automated detection, classification of cell-types and assessing cellular microenvironments in CODEX data with VTEA. Multiplexed immunofluorescence image dataset of human reference kidney was processed, segmented, and analyzed with VTEA. **A**. Maximum projects of subsets of channels highlighting tubular epithelium or immune cells in the renal cortex. Left panel indicates three regions given in **Aa, Ab** and **Ac** and segmentation mask (inset). Scale bars are 500 um and 30 um. **B**. Segmented cells with associated marker intensity with clustered using hierarchical clustering and projected into *t*-SNE space using the average intensity of associated markers. Putative cell-types as indicated. **C**. Marker intensity for clusters identified in **B**, normalized by marker. Cell types include subclasses of epithelial and leukocytes. T_MEM_ and T_N_ are putative memory and novel T-cells. epithelium identified included proximal tubule (PT) S1, S2 and S3 subsegments, loop of Henle thick ascending limb (TAL) and the distal nephron subsegments distal convoluted tubule (DCT) and collecting duct (CD). Markers are given at bottom. A subset of clusters had overlapping proximal tubule (LRP2-positive green arrowhead) and TAL (cyan asterisk) or leukocyte signatures (green and orange asterisks). **D**. Using cell-types defined in **C** and **Figure S3**, neighborhoods were defined in VTEA for every cell within 50 um. The cell census for all neighborhoods was used to cluster and map the neighborhoods to *t*-SNE space in VTEA. **E**. The distribution of cell-types was plotted by neighborhood as a heatmap to identify unique microenvironments in the tissue volume.

## Discussion

In this work, we present an integrated tissue cytometry approach with VTEA to analyze and extract biologically relevant data from state-of-the-art and increasingly multiplexed fluorescence imaging datasets of human kidney tissue. This approach leverages innovative tools for analysis and visualization using machine learning to perform rigorous, reproducible, and informative analysis that could be used to uncover the complex spatial organization and cellular make-up of the human kidney. Using this analysis pipeline, we demonstrated how we can improve the accuracy and resolution of cell classification in kidney tissue. Furthermore, we showed unique advantages of this approach in performing advanced quantitative analysis to uncover cell populations based on spatial associations and neighborhood memberships. In addition, VTEA has the tools to perform intuitive analysis on highly-multiplexed datasets and extract novel information on cell subtypes and neighborhoods that could complement findings from other omics studies at the single cell level. VTEA is available for download through the FIJI plugin updater with the source code is available on github, https://github.com/icbm-iupui/volumetric-tissue-exploration-analysis. Additional description and vignettes demonstrating the use of VTEA are available at https://vtea.wiki.

One of the advantages of incorporating machine learning and dimensionality reduction in the VTEA workflow, as compared to relying only on one label of interest to identify cells, is the ability to use information from the other label intensities and potentially additional spatial parameters. These added parameters increase the discriminative power to identify specific population of cells such as better identification of low intensities of a specific marker (as shown for AQP2). These applications will improve the accuracy compared to a standard gating strategy and increase the confidence of identifying novel cells that may be biologically relevant, as we showed for low AQP2 expressing cells in the distal nephron.

In addition to improving the accuracy and resolution of cell classification, the workflow of VTEA with machine learning tools will facilitate the cytometry analysis of imaging data that was not previously feasible using a standard VTEA approach. We present examples of spatially overlapping cells within structures like glomeruli. Taking advantages of multiple dimensions, it is possible now to accurately quantify the various cell types within glomeruli. This process could also be semi-automated using the data analytical tools provided, thereby having important potential implications on studying human glomerular pathology, where the cell density of specific cell types such as podocyte, immune and mesangial cells may be linked to the pathogenesis of kidney disease^5^. We also demonstrated the utility of VTEA in classifying cells based on spatial parameters, such as neighborhood memberships based on association with structures such as vessels and or other cell types. In the CD45-positive immune cell example, we could classify two cell populations based on nuclear staining and association with vessels. We hypothesize that these immune cell subpopulations may reflect different stages of activity and infiltration: from margination, extravasation and exiting vessels towards forming foci of inflammation within the peritubular space. These findings are only proof-of-principle of the capabilities of VTEA in using imaging-based data for discovery of spatially based cell niches and require further validation to fully determine biological relevance and generalizability^4,19^.

Next, we demonstrated VTEA’s utility in segmentation and analysis of imaging data from kidney tissue, while supporting classification, quantitation, and visualization. VTEA can process mesoscale datasets with tens to hundreds of thousands of cells both in 2D and 3D while maintaining the interactive characteristics of the analysis. In the multiplexed 3D confocal large-scale data, we used unsupervised analysis and dimensionality reduction to classify the cell types and validated these classes of cells based on visualizing the distribution of intensity for each classified cell type. This provides a semiautomated process for large and high-content datasets that augments the rigor of other quality check measures already used, such the validation of the identified cells by mapping them in the original image.

We also analyzed highly multiplexed large-scale data from human cortical kidney tissue imaged with CODEX. This imaging technology expands the ability to multiplex markers on the same 10 µm thick tissue sections using DNA-conjugated antibodies. With CODEX imaging, these antibodies are revealed three at a time by the reversible binding of fluorescent oligonucleotide reporters. Following imaging, the fluorescent reporters are stripped from the tissue and replaced with a second set of three probes and imaged again. This process is repeated until all the antibodies in the tissue have been revealed. Images of DAPI-labeled nuclei are collected in each round to enable registration of images into a single highly multiplexed image. Although CODEX imaging has been described on mouse and human kidney tissue recently^6^, the analytical output from such data has been limited. Using the integrated analytical pipeline with VTEA, we can perform not just cytometry, but also use unsupervised machine learning to classify major cell types and uncover novel subtypes. For example, we demonstrate the existence of subtypes of proximal tubules (PT: CD90-positive PT, PROM1-positive PT) and thick ascending limb cells (PROM1-positive TAL)^4^. Subclassifying also identified T-cell subtypes based on multiple markers. Using neighborhood analysis, we can uncover new spatial associations that could inform on the biology. For example, macrophage association with PROM1-positive TAL cells is consistent with recent single cell transcriptomics data suggesting the transcriptional phenotype of PROM1-positive TALs, may be in niches of immune activation^4^. We expect that applying VTEA analysis on such highly multiplexed CODEX data will complement and spatially anchor single cell transcriptomics data and may inform and confirm (at the protein level within the tissue) transcriptomics outputs such as receptor-ligand analyses.

The advantages of VTEA analysis have been outlined in this work and include the unique integrated workflow in the setting of a general framework of accessibility, flexibility, and extensibility. VTEA can work as a stand-alone tool carrying the imaging data (after collection) all the way to analysis, which offers unique advantages. For example, applying the integrated VTEA workflow to analyze human kidney multiplexed imaging data will enhance efficiency and discovery because all the steps, including advanced machine learning analysis and visualization, occur in one software space. Furthermore, VTEA has unique strengths such as: 1) fine-tuning, in real-time, image processing and segmentation parameters to optimize the analysis and 2) gating for interactive back-and-forth between image and analysis. Importantly, VTEA can also operate with custom workflows to accept inputs from other sources. For example, a better segmentation algorithm (by importing segmentation maps) or an outside set of measurements (such as from MorphoLibJ) can be easily imported. VTEA also exports csv files of measured and unsupervised machine learning features and indexed segmentation maps for integration into other tools and pipelines.

Tissue cytometry with VTEA still has limitations. Despite the advantages of an integrated workflow that has been applied on computer desktops (not requiring computer clusters or server-based), the analysis and visualization of large mesoscale datasets will require desktop computers with enhanced data processing and RAM. Although the analyses of the smaller image datasets presented here can be performed on very modest computers (e.g., 2016 Macbook). There are also persistent challenges in mobilizing large datasets from acquisition platforms to the analysis computers. Importantly, this is not unique to VTEA or even TC and a problem that pertains to other mesoscale and omics datasets. Combining datasets, into the same analytical and non-overlapping image spaces within VTEA is an ongoing area of research and development. Practically, with progress in multiscale and hierarchical image formats this may be greatly simplified soon^32^. Furthermore, multiple datasets can be aligned into a single analysis with VTEA using normalization and mapping strategies but testing and adequately addressing batch effects needs to be established across image datasets^33^. One Possibility is to use cell-centric neighborhood analysis, which is, by default, normalized spatially for a common set of cell classifications, to combine disparate imaging datasets. We recently used such an approach on 3D large-scale multiplex confocal image volumes of human biopsies from the Kidney Precision Medicine Consortium and successfully combined neighborhoods of ∼1.2 million cells^4^.

In conclusion, we present tissue cytometry with VTEA as a solution to analyze and interpret high-content imaging data of human kidney tissue. Using appropriate unsupervised machine learning approaches, we demonstrated how VTEA can classify and characterize cell populations based on a suite of cell-wise features including intensity measurements and neighborhood cell population statistics. We anticipate that this approach will be useful in uncovering the complex spatial organization and cellular make-up of the human kidney and generalized to analyzing imaging data of tissues from other organs.

## Acknowledgments

Imaging was performed in the Indiana Center for Biological Microscopy in the Division of Nephrology, Department of Medicine, Indiana University School of Medicine. We would like to thank Dr. Malgorzata M. Kamocka for assistance with confocal microscopy at the Indiana Center for Biological Microscopy. This project was funded by grants from the National Institute of Diabetes and Digestive Kidney Diseases (NIDDK): PO1DK056788 (TME and JW), P30DK079312 (NIDDK Indiana University O’Brien Center for advanced microscopic analysis-TME), the NIDDK Diabetic Complications Consortium (grants DK076169 and DK115255-TME), and R01DK111651 (TME).

---

**Figure S1.**
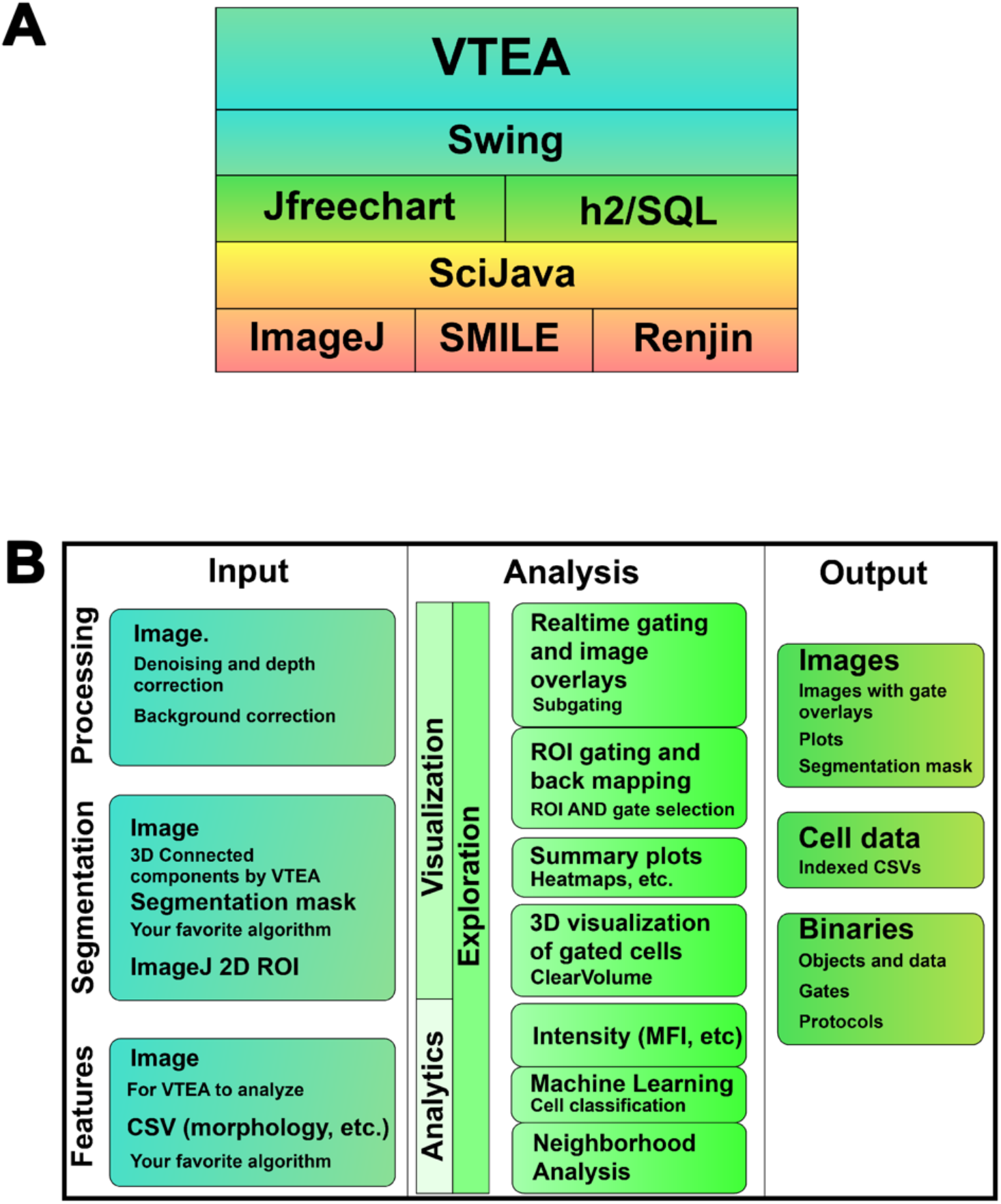
Volumetric Tissue Exploration and Analysis software stack and design schematic. **A**. Volumetric Tissue Exploration and Analysis (VTEA) is built on robust open-source Java projects for visualization, SQL database backend and an annotation-based extensible framework with SciJava for implementing functionality such as image processing (ImageJ), machine learning (SMILE) and the R statistical environment (Renjin). **B**. VTEA is flexible in the data it can operate on (Input, left column), extensible analysis methods it has implement and could be added (Analysis, middle column) and the output data it generates (Output, right column). The output results support management and distribution of data and analysis for scientific rigor and mechanisms for sharing analytical results with other tools such as machine learning pipelines.

**Figure S2.**
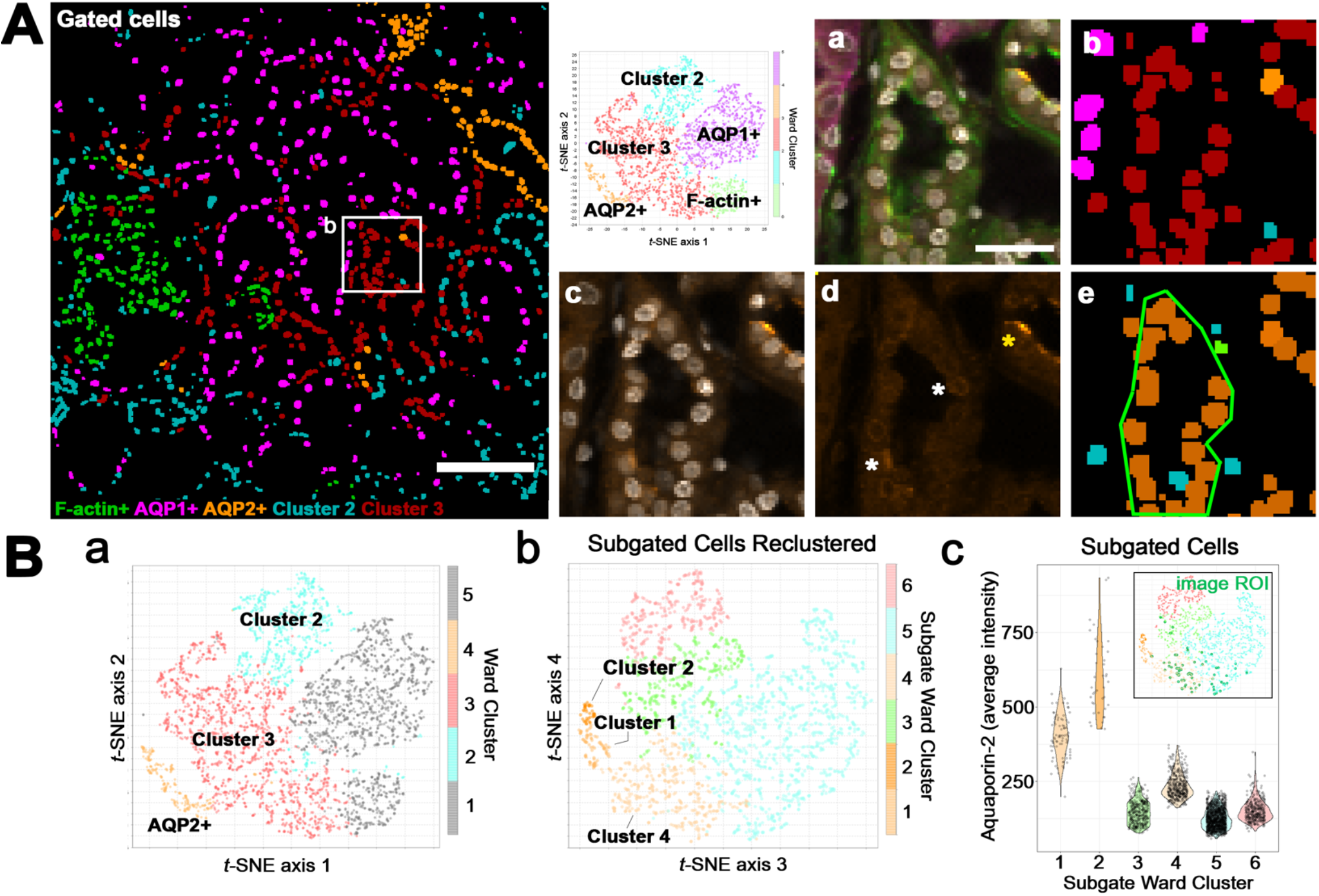
Uncovering subpopulations of AQP2-positive cells with subgating and unsupervised analysis of cell image volumes. The confocal image volume collected and analyzed in Figure 1 was further analyzed for AQP2-positive cells and subpopulations. **A**. Overlays of cell classification as calculated in Figure 1. Low levels of AQP2 are seen in some cells of cluster 3 (inset, **a-d**). Panel **Aa** is an overlay of all four fluorescence images, DAPI in gray, F-actin in green, AQP1 in magenta and AQP2 in yellow. **Ac** and **Ad** show either AQP2 and DAPI or AQP2 alone in the same colors. **Ab** show the overlay of clusters identified in Figure 1, colored by the *t-*SNE given at left. Scale bar = 100 um, A, or 30 um Aa. **B**. Clusters 2-4, low in AQP1 and F-actin intensity (**Ba**), were subgated and clustered and mapped to *t-*SNE space based on the average intensity of all channels for each segmented cell. 3 AQP-positive clusters were identified including the previously mapped collecting duct and a population of AQP2 low epithelium, **Bb** and **Bc**. This was confirmed by image gating and on the cells of interest and mapping to the scatter plot with VTEA (green region-of-interest **Ae** and green rings on scatter plot insert **Bc**). Cluster 1 cells mapped to cells with lower AQP2 intensity, (white asterisks **Ad**) and cluster 2 mapped to cells with higher AQP2 intensity (yellow asterisk **Ad**).

**Figure S3.**
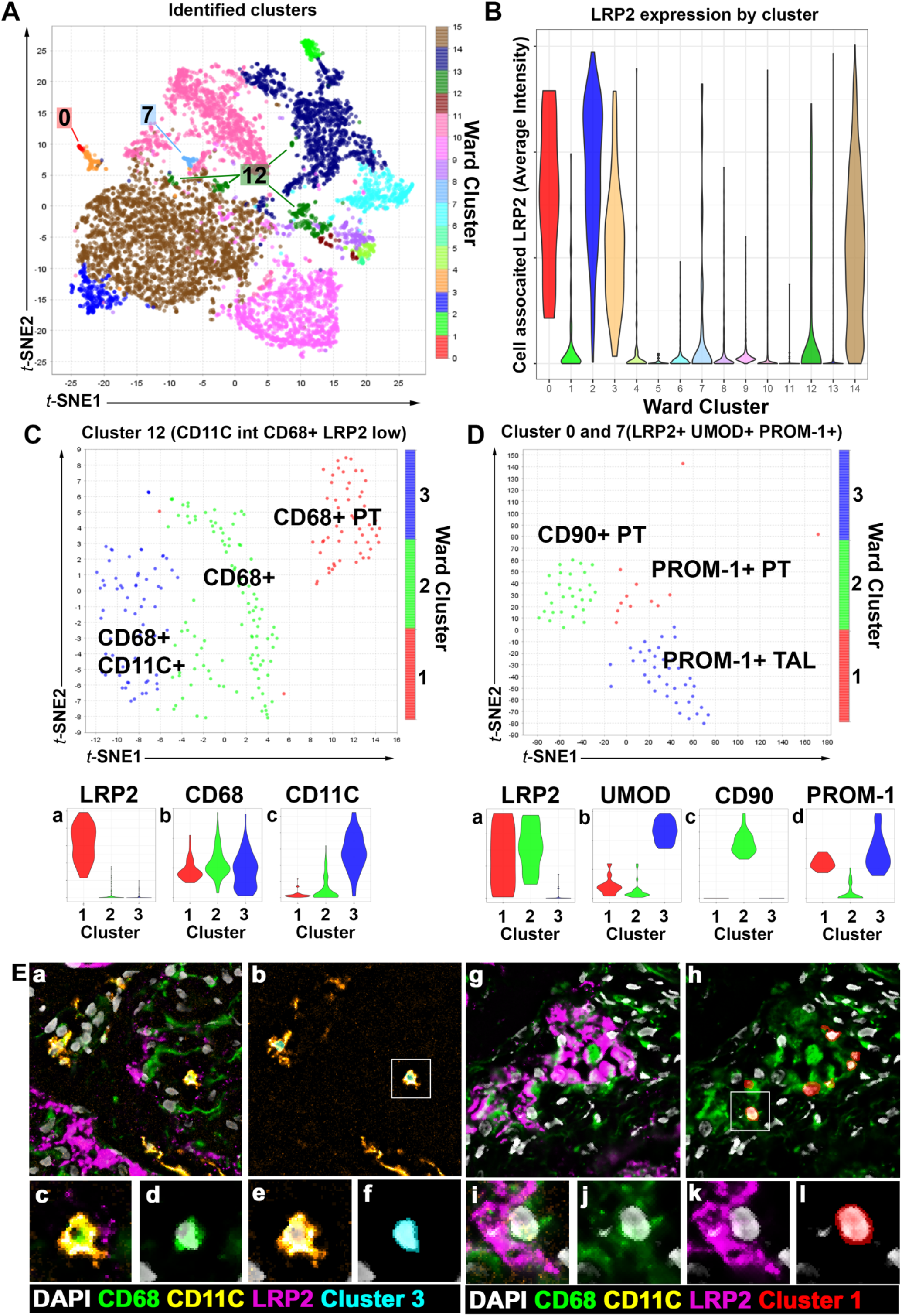
Subclustering of epithelial cells in CODEX data uncovers novel cell states in the proximal tubular (PT) and thick ascending limb (TAL). **A-B**. Clusters 12 or clusters 0 and 7 from Figure 7 were subgated based on intermediate LRP2 expression and either CD68 or PROM-1 or UMOD expression. **C**. Cluster 12 was reclustered separating CD68+ DCs, putative macrophages and CD68-positive PT cells. **D**. Clusters 0 and 7 were subgated and reclustered identifying two PT cell-types (PROM-1-positive vs. CD90+). **E**. CD68-positive DC-cells and putative epithelium are readily identifiable (**Ea-f** vs **Eg-m**). **Ec-f** and **Ei-l** are insets for **Ea** and **Eh** respectively.

